# Linkage Disequilibrium and Inversion-Typing of the *Drosophila melanogaster* Genome Reference Panel

**DOI:** 10.1101/014936

**Authors:** David Houle, Eladio J. Márquez

## Abstract

We calculated the linkage disequilibrium between all pairs of variants in the Drosophila Genome Reference Panel with minor allele count equal to or greater than 5. We used *r*^*2*^≥0.5 as the cutoff for a highly correlated SNP. We make available the list of all highly correlated SNPs for use in association studies. Seventy-six percent of variant SNPs are highly correlated with at least one other SNP, and the mean number of highly correlated SNPs per variant over the whole genome is 83.9. Disequilibrium between distant SNPs is also common when minor allele frequency (MAF) is low: 37% of SNPs with MAF<0.1 are highly correlated with SNPs more than 100kb distant. While SNPs within regions with polymorphic inversions are highly correlated with somewhat larger numbers of SNPs, and these correlated SNPs are on average farther away, the probability that a SNP in such regions is highly correlated with at least one other SNP is very similar to SNPs outside inversions. Previous karyotyping of the DGRP lines has been inconsistent, and we used LD and genotype to investigate these discrepancies. When previous studies agreed on inversion karyotype, our analysis was almost perfectly concordant with those assignments. In discordant cases, and for inversion heterozygotes, our results suggest errors in two previous analyses, or discordance between genotype and karyotype. Heterozygosities of chromosome arms are in many cases surprisingly highly correlated, suggesting strong epsistatic selection during the inbreeding and maintenance of the DGRP lines.

## INTRODUCTION

The Drosophila Genome Reference Panel (DGRP; Mackay et al. 2012) is a set of sequenced inbred lines derived from a single outbred population of *Drosophila melanogaster*. The DGRP has been used for a series of genome-wide association (GWA) studies on a wide variety of phenotypes. Linkage (gametic-phase) disequilibrium (LD) is a challenge to all GWA studies, as it confounds the signal from variant sites (we call these SNPs for brevity) that cause phenotypic variation with those that are genetically correlated with the causal variant but that do not have effects on the phenotype. The nature of the DGRP in many ways minimizes the presence of LD relative to vertebrates or to line-cross-derived mapping populations. The DGRP lines are drawn from a natural population in Raleigh North Caroline with large effective size, as shown by the relatively low level of structure within the population. Mackay et al. (2012) confirmed that the average LD drops very rapidly with distance between SNPs, to an average squared correlation r^2^<0.2 at just 10 base pairs on the autosomes. This result might suggest that the overall impact of LD on GWAS results will be low.

More detailed analyses (Huang et al. 2014; Pool 2015; Cridland et al. 2013) show evidence for potentially troublesome LD within the DGRP. Huang et al. estimated that 2.7% of line pairs have relatedness greater than 5%, and 0.05% of pairs (11 pairs) are related at greater than 50% (c.f. Cridland et al. 2013). A total of 16 alternate chromosomal inversion karyotypes are present in the DGRP (Corbett-Detig and Hartl 2012; Huang et al. 2014; Langley et al. 2012). Huang et al. (2014)’s analysis suggests that three of these are fixed in seven or more lines. These more common inversion types are substantially differentiated from the Standard karyotypes, and cause LD (Corbett-Detig and Hartl 2012; Langley et al. 2012; Huang et al. 2014). Huang et al. also showed that rare SNPs have a substantial likelihood of being highly correlated with SNPs that are more than 1 kb distant even outside of inversions. Finally, Pool (2015) analyzed whether small genomic regions in the DGRP lines were likely to reflect African rather than European ancestry, following the joint colonization of North America by a mixture of *D. melanogaster* from these two source populations (Duchen et al. 2013). He found that about 20% of the DGRP genomes could be assigned African ancestry. In addition there was significant LD between these African regions among different chromosomes, suggesting possible epistatic selection favoring genotypes from the same region.

These findings document the problem of LD, but to interpret associations between SNPs and phenotypes in the DGRP we need to know whether particular SNPs that are implicated are correlated with other SNPs or inversions, and where those correlated sites are. We calculated the LD between all pairs of SNPs in the 205 Freeze2 DGRP lines, and provide a comprehensive list of polymorphic sites in substantial LD with inversions and with other sites throughout the genome. The cytogenetic karyotype assignments in Huang et al. (2014) do not always agree with other PCR-based or sequence-based assignments in two other papers (Corbett-Detig and Hartl 2012; Langley et al. 2012), and we use genotypic data to investigate why these assignments differ.

## METHODS

We used Freeze 2 genotype calls for the DGRP lines obtained from ftp://ftp.hgsc.bcm.edu/DGRP/freeze2_Feb_2013/freeze.vcf. We used only calls with genotypic phred scores ≥20 at sites with exactly two alternative types. For LD calculations, heterozygous calls were treated as missing data. We focused our attention on linkage disequilibrium at focal sites with minor allele counts (MAC) of 5 or more, although we calculated correlations of these focal sites with those where MAC was three or more. We excluded sites where the number of missing calls was greater than 85. There are 2,659,276 focal sites with MAC≥5, and 3,159,155 sites with MAC≥3. For focal sites, the median minor allele frequency was MAF=0.13, and the median number of lines scored was 195 out of 205 possible.

We parameterized linkage disequilibrium (LD) as the product-moment correlation *r*^2^ (Hill and Robertson 1966). Our algorithm for finding such pairs is based on the fact that only SNPs with similar minor allele frequencies (MAF) could be highly correlated. The *r*^2^ value used as a cutoff dictates the degree of similarity possible; for rare SNPs, only sites with *r*^*2*^*p* <MAF<*p/r^2^* can be correlated to that degree with the focal SNP. For reasons discussed below, we chose to use *r*^2^≥0.5 as our cutoff for LD. In general, the relationship between MAF and the maximum correlation is non-linear, so we calculated an empirical estimate of how similar MAF values could be and yield *r*^2^≥0.5. Virtually every site had some missing calls, and the overlap of those missing calls is different for different sites necessitating an approximate solution to this limit problem. We first binned SNPs into frequency classes to the nearest 0.01. We then fit a quadratic equation to the upper limit of MAF that could be correlated at *r*^2^>0.5 for SNPs at the upper limit of the bin.

To calculate LD between all pairs of SNPs within this limit, we started with the lowest frequency bin, whose rounded frequency is *p*_*b*_, and calculated correlations between all SNPs in the focal bin and those SNPs with MAF below the empirically determined limit *p*_*b*_ + 0.005 < MAF < 0.01 + 1.875*p*_*b*_ − 1.17 (*p*_*b*_)^2^. This process was repeated for each bin with larger MAF. We retained a list of all pairs of SNPs with *r*^*2*^>0.5. SAS programs to calculate the MAF limits for a given **r*^*2*^* cutoff, and for calculating and storing the identity of SNP pairs with high *r*^*2*^ are included in Supporting Information, File S1. This algorithm will miss a small number of highly correlated SNPs that have missing genotype information in many lines.

Prior to carrying out the above calculation, we characterized the probability that a SNP correlated with a causal SNP will result in a false positive using simulated data. We simulated phenotypic effects on a set of correlated SNP genotypes drawn from the Freeze1 genotype calls for 165 DGRP lines (Mackay et al. 2012; ftp://ftp.hgsc.bcm.edu/DGRP/freeze1_July_2010/snp_calls/). We first identified all SNP pairs correlated at *r*^2^>0.25 using an algorithm similar to that outlined for the Freeze 2 data, outlined above. We then drew 100 random focal SNPs that were each correlated at *r*^2^>0.25 with at least one other SNP, which we call a SNP family. Ten focal SNPs were chosen from each MAF decile. In cases where the focal SNP was correlated at *r*^2^>0.25 with more than 100 SNPs (31% of all focal SNPs), we retained a random sample of 100 correlated SNPs. We simulated SNP phenotypic effects that explained 1% of the total phenotypic variance in a multivariate trait. Simulated data were analyzed using MANOVA, with SNP genotype as the sole predictor variable. MANOVA P-values were calculated using a chi-square approximation of Wilk’s Lambda.

Huang et al. (2014) reported that three rare inversion karyotypes were fixed in seven or more DGRP lines (In(2L)t, In(2R)NS, In(3R)Mo), while no other karyotype was fixed in more than four lines. These karyotype assignments are sometimes in disagreement with the sequence-based assignments of karyotype reported by Corbett-Detig and Hartl (2012) and Langley et al. (2012). We checked these characterizations statistically using the following approach. We assembled genotypic data from Freeze 2 as above, but including heterozygous assignments, then excluded SNPs with five or more missing genotype assignments. Missing assignments in the remaining SNPs were assigned to the common allele to provide complete genotypic data. Using the results from the genome-wide LD results, we then obtained a list of the SNPs that are inside the inversion breakpoints of the three common alternative karyotypes (Corbett-Detig and Hartl 2012; Corbett-Detig et al. 2012), and that had LD *r*^*2*^>0.5 with 200 or more other SNPs more than 100k sites distant. These SNPs are likely to be characteristic of inversion types. This provided a sample of 28,495 SNPs on chromosome 2L, 8,174 on 2R and 8,347 on 3R. We conducted separate principal components analyses (PCA) of the genotypes of a randomly chosen subset of 5,000 SNPs for each inversion, and used the scores on PC1 to diagnose which genotypes are characteristic of each karyotype. We also calculated the proportion of SNPs that were scored as heterozygous for chromosomal regions defined by the inversion breakpoints.

The African ancestry of each inverted region was calculated using the results of Pool (2015). His Table S3 lists regions identified as having a high probability of African ancestry based on applying the Hidden Markov Model of ancestry from Pool et al. (2012) to the DGRP lines. We calculated the lengths of these regions that lay between the breakpoints of the three common inversions for each DGRP line.

Calculations were carried out in SAS version 9.3 for Windows and Unix (SAS Institute 2011).

## RESULTS

To characterize linkage disequilibrium (LD) in Freeze 2 of the DGRP, we considered all pairs of sites with minor allele counts (MAC) ≥5. These sites include indel variation, but we will refer to them as SNPs for brevity. The overall magnitude of linkage disequilibrium (LD) in Freeze 2 of the DGRP for 1 million pairs of random SNPs with MAC ≥5 is shown in Figure 1A. We use *r*^2^ as our measure of disequilibrium (Hill and Robertson 1966) because this is the most appropriate indicator of the likelihood that analyses of pairs of SNPs will yield similar results. Over 99.9% of all random SNP pairs are correlated at less than *r*^*2*^≤0.15, and only 0.0024% have *r*^*2*^≥0.5. There are however, almost 3.5×10^12^ SNP pairs for this data set, so the number of pairs correlated at any particular level is not small.

**Figure 1.**
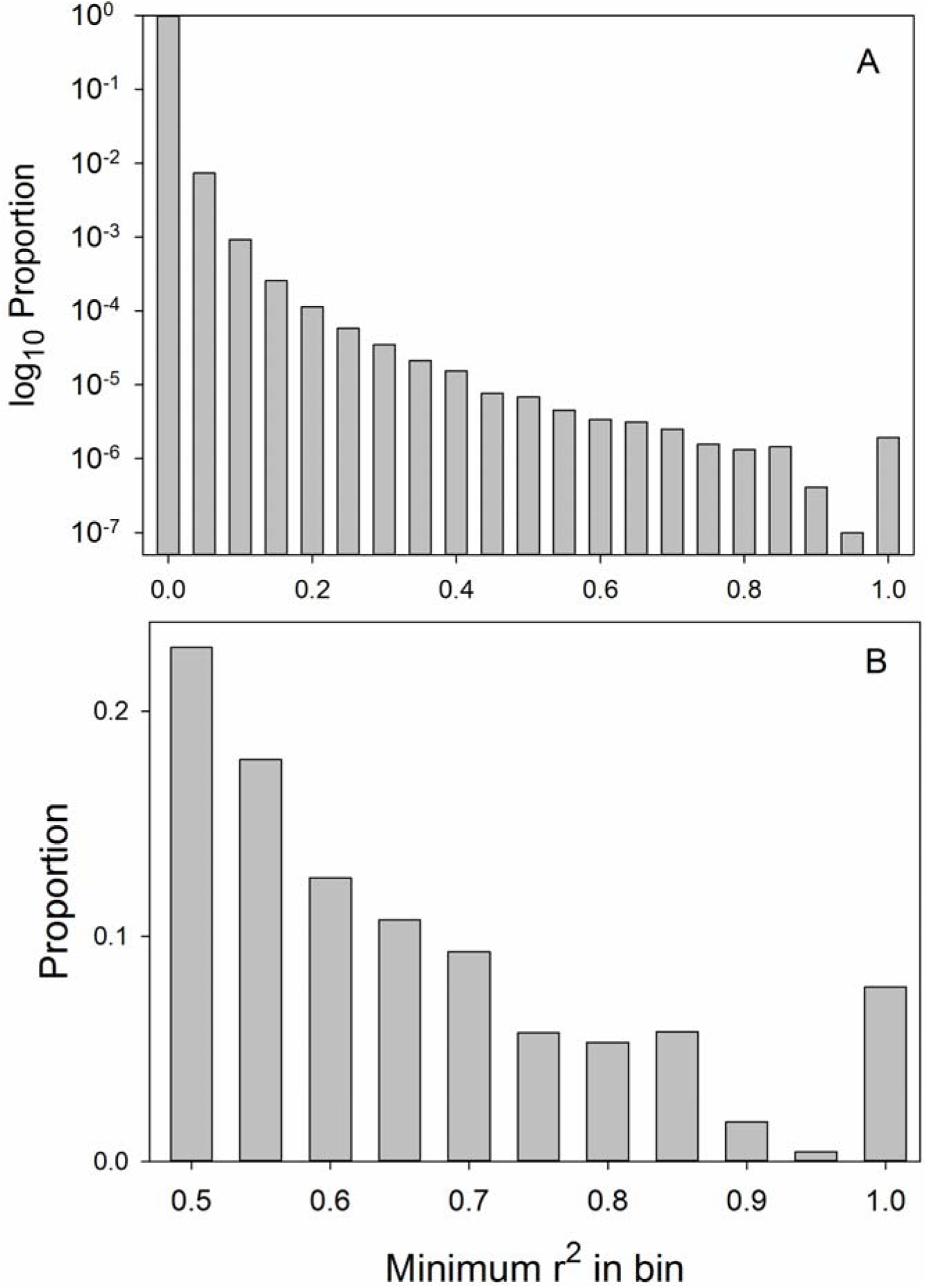
Distribution of LD between pairs of SNPs, measured as r^2^. *A*. Distribution of LD among 1,000,000 random SNP pairs. Note the log_10_ scale. *B*. Distribution of r^2^ values among all highly correlated pairs with MAC≥5 genome-wide.

To determine what level of *r*^*2*^ we should focus on, we carried out simulations in which we simulated a phenotypic effect at a focal SNP, then assessed the probability that correlated SNPs that do not themselves cause phenotypic variation would show a significant phenotypic effect, with the results shown in Figure 2. We simulated effect sizes to generate modest power of about 0.3 at a conservative *P* value of 10^-6^, likely to be typical of many studies using the DGRP lines. Fig. 2 shows that, as long as *P* values are not very liberal, *r*^*2*^≥0.5 will include the vast majority of SNPs likely to generate false positives due to LD. Consequently, we structured our calculations to ensure that the vast majority of correlations of *r*^*2*^≥0.5 would be detected, as described in Methods.

**Figure 2.**
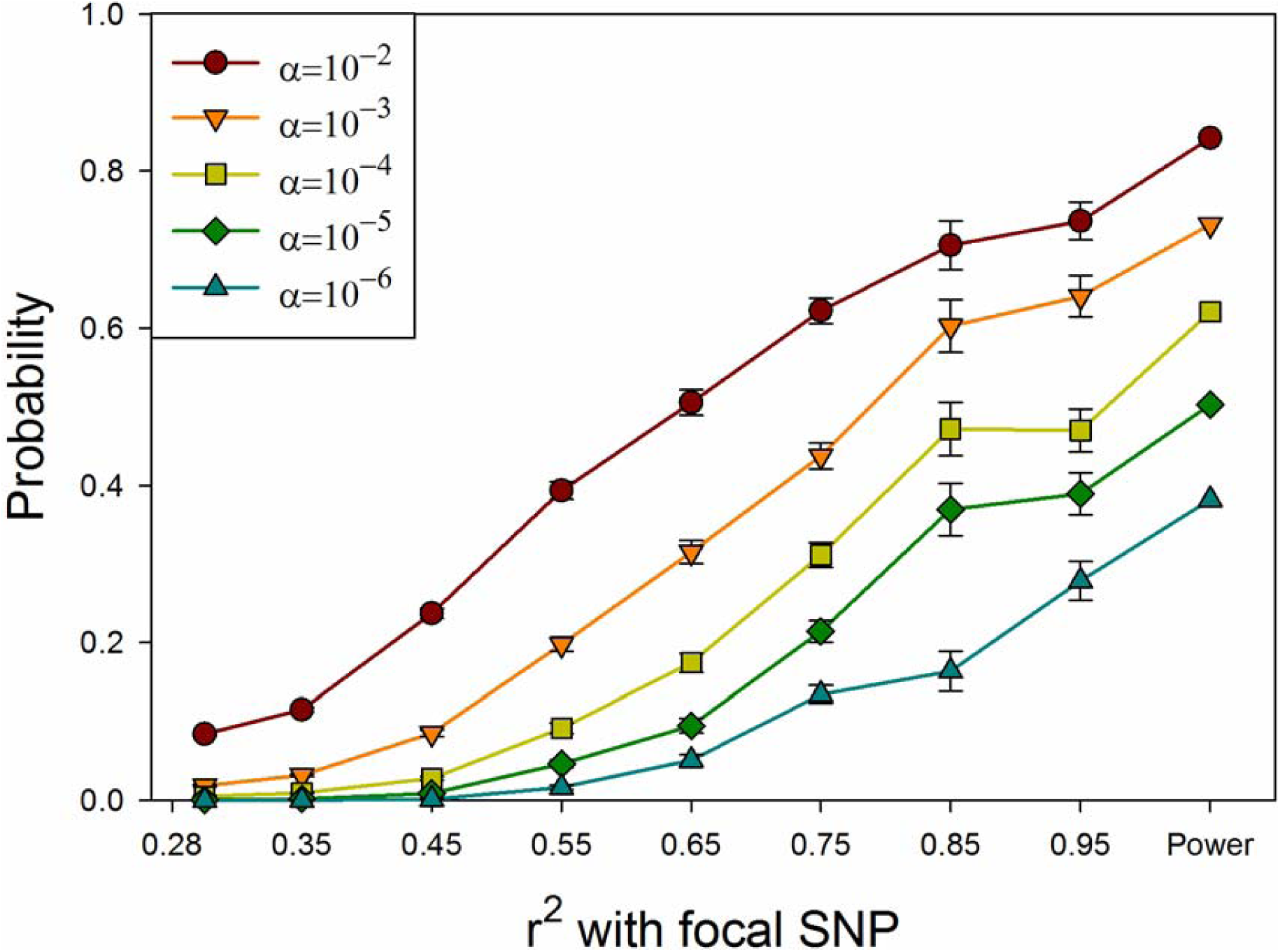
Probability that a SNP with no phenotypic effect will have a statistically significant association with phenotype as a function of LD with a SNP that does have a phenotypic effect. α= threshold for significance. The focal SNP explains 1% of the phenotypic variance in a multivariate trait. The rightmost symbol is power to detect an association with the focal SNP. Values on the X-axis are the midpoints of bins of r^2^ values.

**Figure 3.**
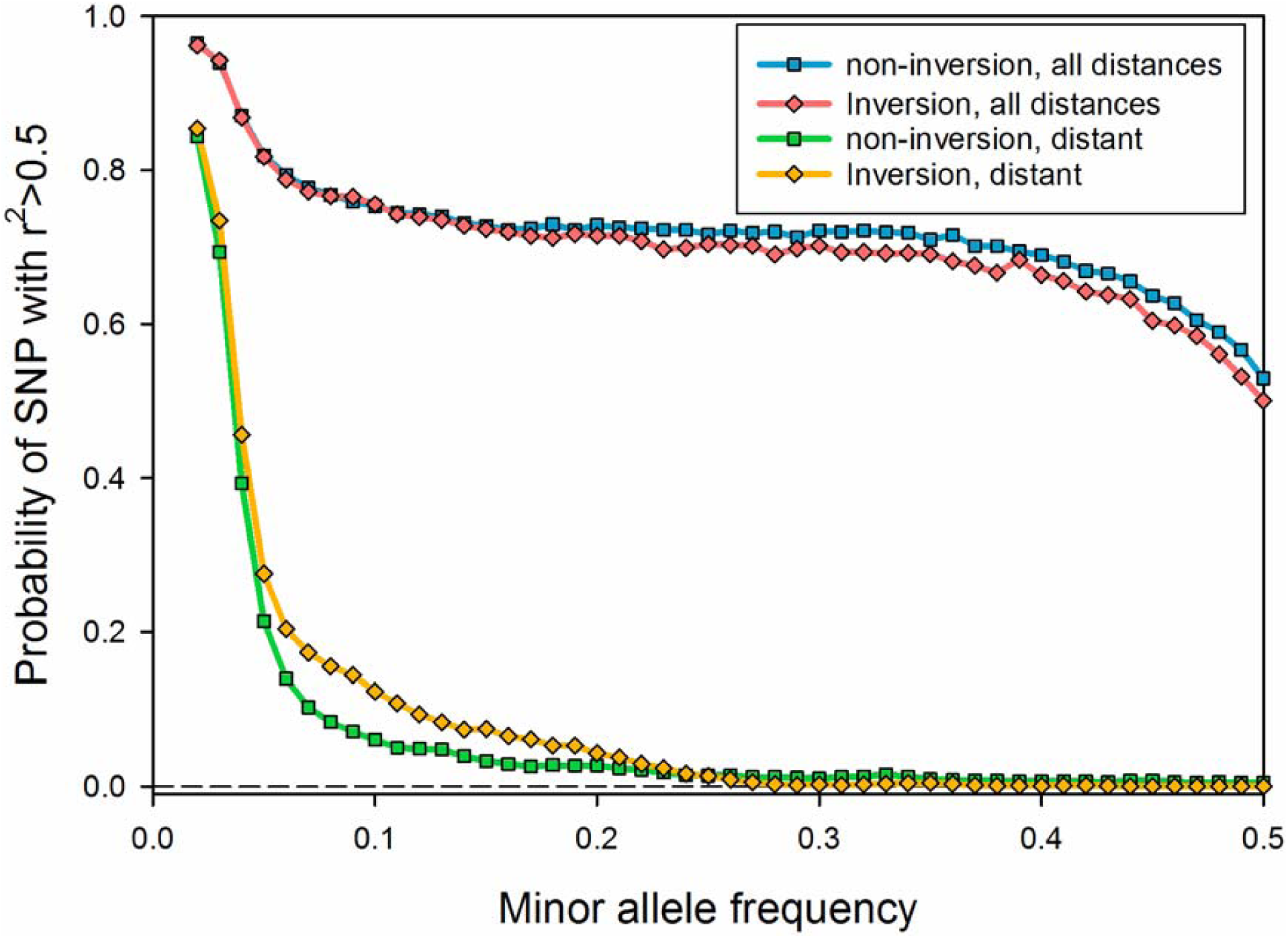
Probability that a focal site is correlated at r^2^>0.5 with at least one other site in the genome, as a function of location relative to inversions. Inversion sites lie between the breakpoints defined by Corbett-Detig et al. (2012), with the exception of the distal segment of chromosome 3R, which was treated as part of In(3R)Mo (Corbett-Detig and Hartl 2012).

We refer to pairs of SNPs correlated at *r*^*2*^*≥*0.5 as “highly correlated,” and SNPs more than 100kb apart as “distant”. To provide a comprehensive picture of LD for these SNPs, we also calculated LD with SNPs that had a MAC as low as three. A total of 1.36×10^8^ highly correlated pairs were detected. More importantly, 76% of all SNPs with MAC ≥5 were highly correlated with at least one other SNP. Seventeen percent of all SNPs are highly correlated with at least one distant SNP, and 7.7% are highly correlated with a SNP on another chromosome. The distribution of *r*^*2*^ values for highly correlated pairs where both have MAC ≥5 is shown in Fig. 1B. Due to the predominance of rare minor alleles (Mackay et al. 2012), 38% of all high correlations for sites with MAC ≥5 involve sites with MAC<5. We did not include these in Fig. 1, but they have a strong mode around *r*^2^ ≈ 0.6 due to SNPs that match at all but one site. The full lists of highly correlated SNPs (including those between sites with MAC ≥5 and sites with MAC<5) by chromosome arm are available in Supporting Information, File S2.

Fig. 3 shows the probability that a SNP is highly correlated with at least one other SNP. SNPs were classified as inside or outside the breakpoint of the three inversions fixed in more than four DGRP lines (In(2L)t, In(2R)NS, and In(3R)Mo). For local disequilibrium, it makes little difference whether a SNP is in an inversion or not, but SNPs in inversions are more likely to be in high LD with a SNP that is distant (in the Standard gene order). The inversion karyotypes themselves have frequencies of 0.1 or less (see below), precluding SNPs characteristic of inversions from being highly correlated with high MAF variants. More importantly, simply excluding SNPs within inversions does not appreciably reduce the likelihood that a variant will be highly correlated with at least some other SNPs, nor preclude those highly correlated SNPs from being distant.

Figure 4 shows the mean and median numbers of SNPs highly correlated with a focal SNP for all pairs of SNPs, and all distant pairs. The numbers of highly correlated pairs are substantially higher at low MAF. The difference is particularly large for distant SNPs. When MAF<0.1, 37% of SNPs are correlated with a distant variant; when MAF is between 0.1 and 0.2, 5% of SNPs are. On the other hand, the mean number of highly correlated SNPs is still substantial at all allele frequencies. This is consistent with random disequilibrium due to the very large number of SNPs with low MAF (Mackay et al. 2012), and to the smaller number of permutations that can lead to a low MAF. We refer to this as rarity disequilibrium. Of the 2.6 million SNPs with MAC≥5 in this analysis, 50% have MAF<0.129, and 25% have MAF<0.054. Supporting Information Figure S1 suggests that inversions may compound the effects of linkage and rarity disequilibrium, as the mean number of highly correlated SNPs inside inversions is substantially higher when MAF is less than 0.2. This is particularly so for SNPs distant from the focal variant.

**Figure 4.**
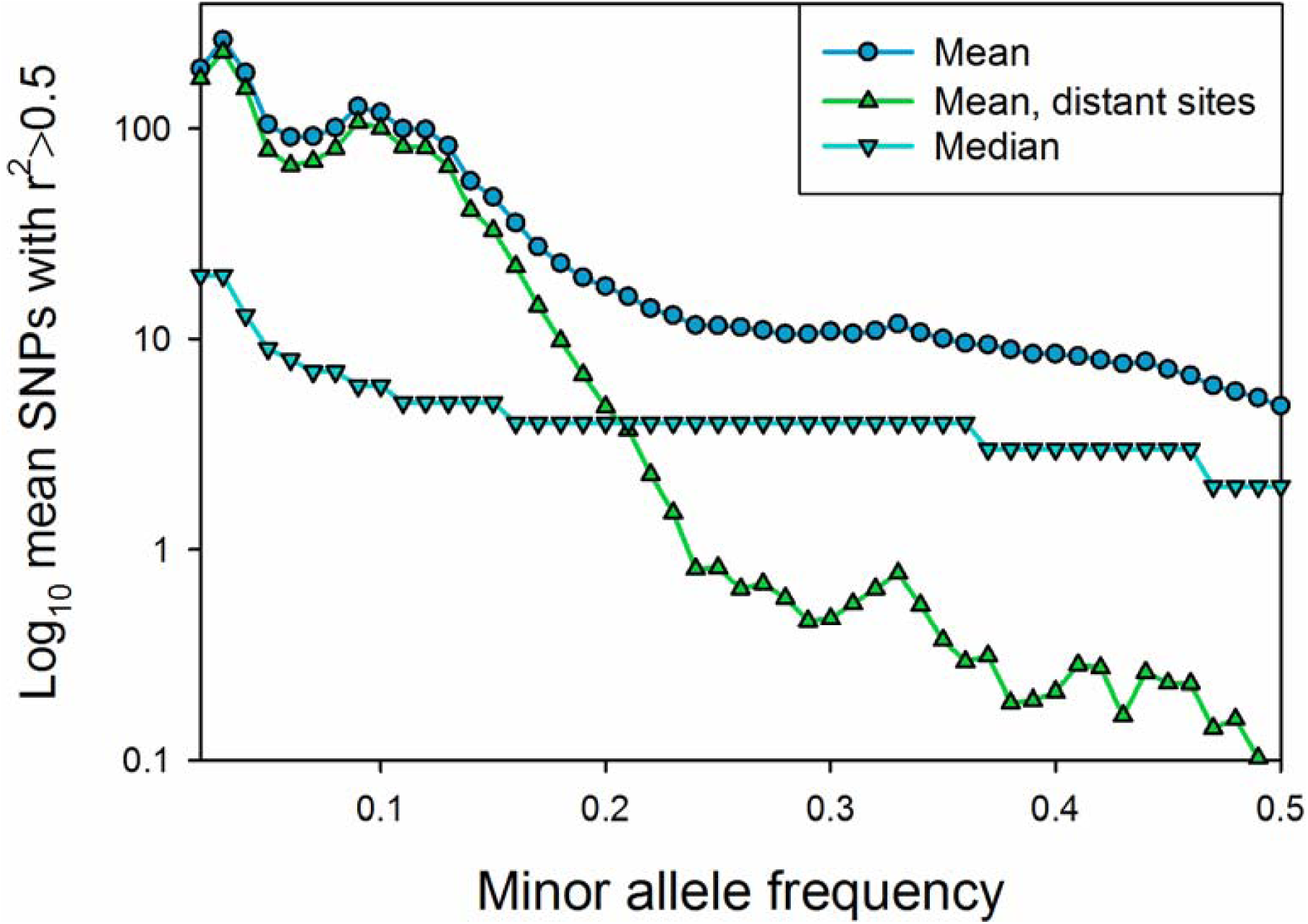
Mean and median number of sites correlated with variant sites at r^2^>0.5 as a function of minor allele frequency.

The variance in the number of correlated SNPs is high and skewed towards smaller numbers, so that the mean number of correlated SNPs is quite a bit higher than the median, as shown in Figure 4. The median number of correlated SNPs across the genome nevertheless ranges from 20 for most of the lowest MAF classes to 2 for the high MAF SNPs. The median number of SNPs greater than 105 bp away that are in high LD is 1 or more when MAF is 0.04 or less, but 0 for all higher frequencies. The median number of variants correlated at *r*^*2*^>0.5 is virtually identical between SNPs inside and outside of inversions, regardless of MAF (not shown).

The mean number of highly correlated distant sites across the genome is shown in Figure 5. There is a clear peak of distant LD in In(2R)NS, and near the breakpoints in In(2L)t. There is also a broader, but less intense peak of distant LD in the neighborhood of In(3R)Mo, but for this arm, high LD extends beyond the proximal and distal breakpoints of this inversion. The high LD between the distal end of 3R and In(3R)Mo has been noted before (Corbett-Detig and Hartl 2012). Other low recombination regions near telomeres and centromeres also show high distant LD.

**Figure 5.**
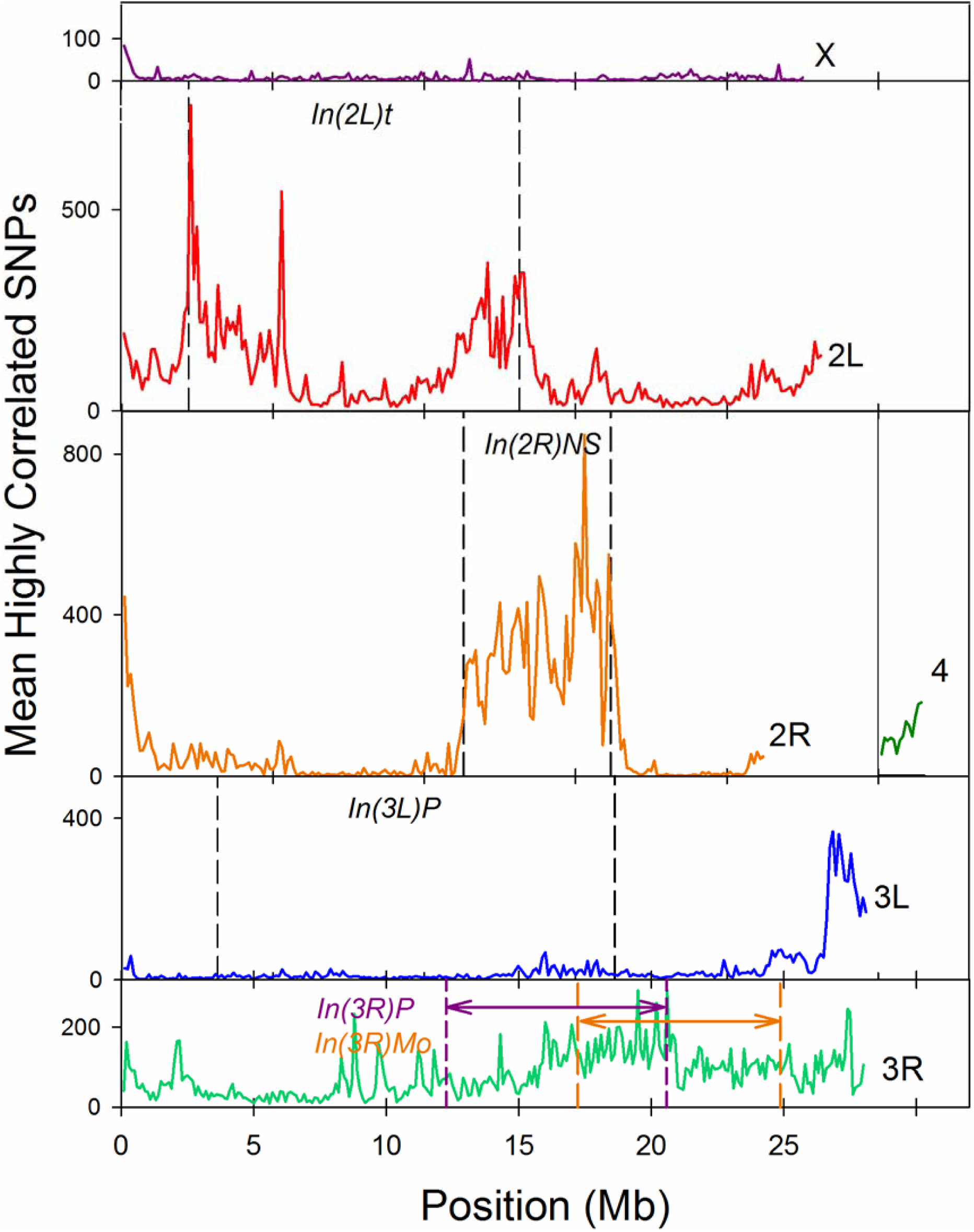
Mean number of highly correlated distant sites/SNPs in 100kb regions across the genome.

The cytogenetic analyses of Huang et al. (2014) found at least seven DGRP lines fixed for each of the common cosmopolitan inversions In(2L)t, In(2R)NS, and In(3R)Mo, and many more lines that were heterozygous for these karyotypes. The remaining inversion karyotypes were fixed in four or fewer lines. Two previous studies (Corbett-Detig and Hartl 2012; Langley et al. 2012) also inversion-typed a subset of the DGRP lines using either a PCR- and/or a next-generation-sequence-based assay. These two studies were consistent in their assignments, so we refer to them collectively as CD-L. CD-L and Huang et al. both scored 501 chromosome arms for homozygotes of the three common inversions In(2L)t, In(2R)NS, and In(3R)Mo. Ten of these assignments are in conflict, and 459 are in agreement. The remaining 32 were scored as heterozygotes by Huang et al. but were not examined by Langley et al. (2012). Corbett-Detig and Hartl (2012) state they did detect heterozygotes for inversions, but only reported lines positively identified as inversion homozygotes.

Given the pattern of distant LD shown in Fig. 5, and because each of these previous studies of inversion type makes clear that inversion karyotypes are usually substantially differentiated from each other, we reasoned that SNPs within the inversion breakpoints that show high LD with a large number of distant SNPs will tend to be diagnostic for inversion type, and identify the likely source of discrepancies between previous karyotype assignments. After selecting SNPs with nearly complete genotypic data that are also in high LD with many other SNPs, we performed principal components analyses on those SNPs located within the breakpoints of each of the three common inversions.

The full list of previous karyotype inferences, scores on the first PC for inversion-diagnostic genotypes for each chromosome, and heterozygosities of all SNPs within regions defined by inversion breakpoints are given in Supporting Information, File S3. We plot the PC1 scores vs. the average heterozygosity (H) of each inversion region in Figure 6. In all but one of the 458 cases where Huang et al. and CD-L both reported a homozygous karyotype, scores on PC1 predict inversion type. The exception is line RAL332 for chromosome 3R, where both Huang et al. and CD-L infer the Standard arrangement, while PC1 score and the observed heterozygosity predict a Standard/In(3R)Mo heterozygote. Intermediate scores on PC1 are found in chromosome arms identified as inversion heterozygotes by Huang et al, with four exceptions. PC1 score for line RAL325, chromosome 2R indicates a Standard/In(2R)NS heterozygote, while Huang et al. reported two different inversions as heterozygotes, In(2R)Y6 and In(2R)Y7. RAL 409 is anomalous in having PC scores suggesting heterozygotes for In(2R)NS and In(3R)Mo, but fairly low heterozygosity. It was scored a homozygote for In(2R)NS and In(3R)Mo by both Huang et al. and CD-L.

**Figure 6.**
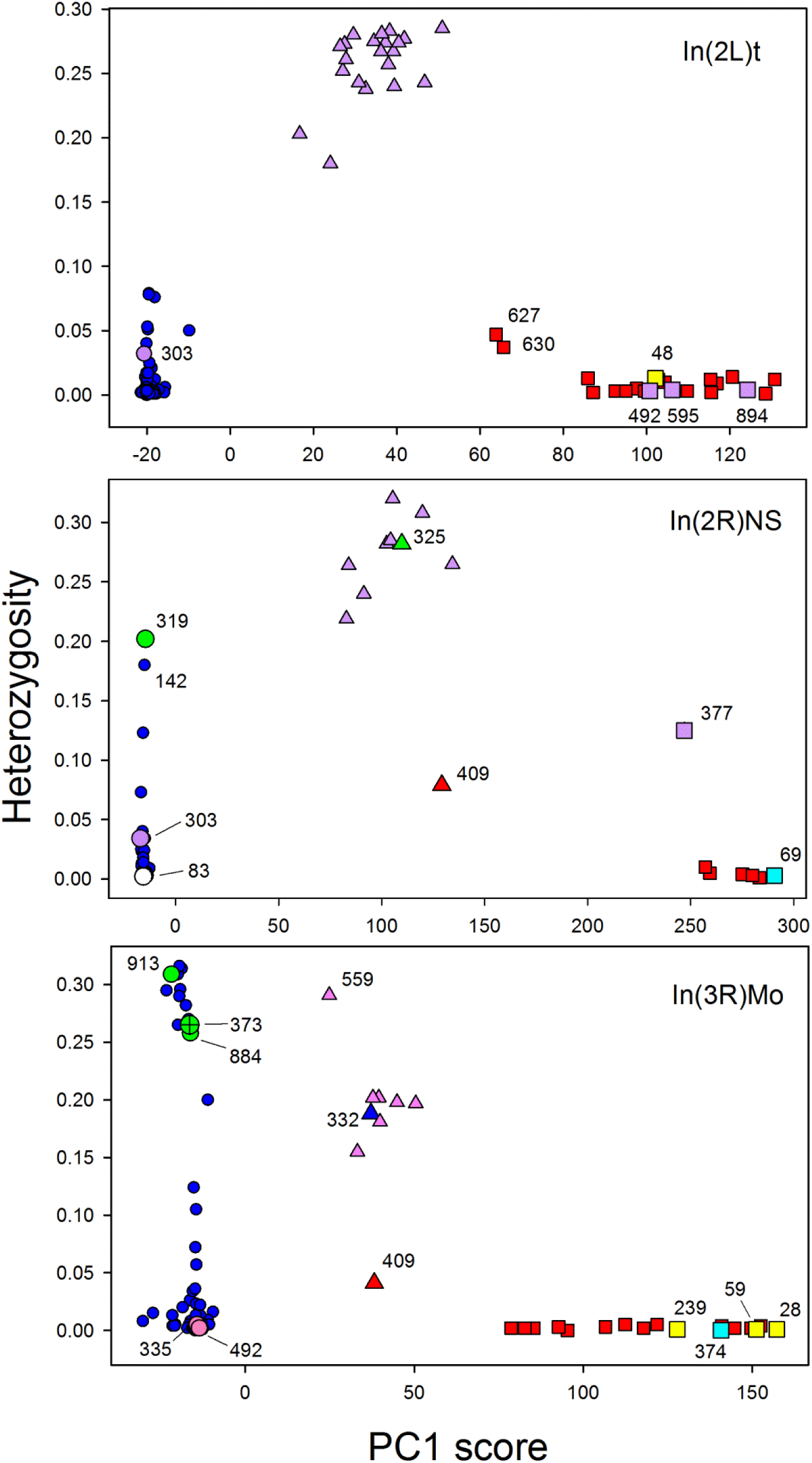
Inferred karyotype and heterozygosity of DGRP lines for common inversions. DGRP line numbers indicated for anomalous cases. Predictions based on our principal components analyses are shown by symbol shapes: squares=inversion homozygotes, circles=Standard homozygotes, triangles=inversion heterozygotes. Colors reference the states inferred in Huang et al. and CD-L: Red=inversion homozygote in both; blue=standard homozygote in both; lavender=heterozygous in Huang et al. (CD-L in most cases did not scores heterozygotes); yellow=Standard homozygote in Huang et al., inversion homozygote in CD-L (exc. RAL48, not scored by CD-L); cyan=inversion homozygote in Huang et al., standard homozygote in CD-L; green=predicted heterozygote for a rare inversion (Line 373 for 3R was predicted as In(3R)Mo homozygote by CD-L); white=Standard homozygote in Huang et al., In(2R)NS homozygote by CD-L. Note that separate PC analyses were carried out separately for each inverted region, so the scale of PC1 scores for each chromosome arm is different.

A number of inversion regions have highly heterozygous sequence data (H>0.15) but no evidence of similarity to common inversion genotypes. In four cases (shown in green in Fig. 6), these were identified as heterozygotes for rare inversion karyotypes by Huang et al. Line RAL303 was scored as an inversion heterozygote for both In(2L)t and In(2R)NS by Huang et al., but does not have a genotype characteristic of either heterozygote. Fourteen lines for which neither PC1 scores nor Huang et al. suggest inversion heterozygosity are highly heterozygous in the region of In(3R)Mo, which may suggest the presence of balanced polymorphism not associated with an inversion.

There are a total of six cases where CD-L and Huang et al. assigned different homozygous karyotypes to the same lines; in three cases PC1 scores are consistent with CD-L, while PC1 scores and Huang et al. are in agreement for the other three. Huang et al. reported an additional five cases of karyotypic heterozygosity that do not have elevated sequence heterozygosity.

Inversions In(2L)t and In(2R)NS both have African origin, while In(3R)Mo has a non-African origin (Corbett-Detig and Hartl 2012). Pool (2015) inferred homozygous regions in each DGRP line that were likely to have African ancestry, and we used the total length of these in each inverted region to provide an additional check on our inversion-typing, with results shown in Supporting Information, Table S1. Mean African ancestry for our predicted inversion types agrees well with expectations for In(2L)t and In(3R)Mo. For In(2R)NS, however, the two mismatches between our predictions and previous studies both have African ancestry more consistent with the opposite type. The highest African ancestry in the 187 consensus Standard karyotypes is 920kb, while line 83 has African identity for 5,840kb. The lowest African ancestry in the 7 consensus In(2R)NS lines is 3,286kb, while line 69 has African identity for just 874kb. These are both cases where the sequence-based typing of Corbett-Detig and Hartl (2012) is in disagreement with our predictions and karyotyping by Huang et al. (2014). These two lines may represent recent double recombinants that separate the In(2R)NS breakpoints from their typical genotype.

Heterozygosities of segments of chromosome arms defined by common inversion breakpoints are correlated, as shown in Table 1. Segments of the same arm always have correlations of 0.86 or above. Segments of different arms of the same chromosome also remain highly correlated. These results suggest that there is strong selection against recombinants and segregants in some of the DGRP lines. It is particularly striking that X-chromosome heterozygosity is significantly correlated with the heterozygosity of chromosome 3.

**Table 1.** Pearson correlations of heterozygosities for regions of chromosome arms.

| chromosome<br>region | In(2L)t | prox. 2L | distal 2R | In(2R)NS | prox. 2R | 3L | prox. 3R | In(3R)Mo | distal 3R | X | H ± S.D. |
| --- | --- | --- | --- | --- | --- | --- | --- | --- | --- | --- | --- |
| distal 2L | 0.93** | 0.92** | 0.45** | 0.40** | 0.66** | 0.03 | -0.01 | -0.01 | -0.08 | 0.10 | 0.031 ± 0.065 |
| In(2L)t | | 0.98** | 0.39** | 0.36** | 0.57** | -0.01 | -0.04 | -0.04 | -0.10 | 0.09 | 0.032 ± 0.031 |
| proximal 2L | | | 0.43** | 0.38** | 0.58** | -0.01 | -0.05 | -0.04 | -0.11 | 0.08 | 0.030 ± 0.077 |
| distal 2R | | | | 0.90** | 0.87** | 0.09 | -0.08 | -0.08 | -0.09 | 0.06 | 0.019 ± 0.054 |
| In(2R)NS | | | | | 0.86** | -0.02 | -0.06 | -0.06 | -0.06 | 0.06 | 0.020 ± 0.060 |
| proximal 2R | | | | | | 0.00 | -0.02 | -0.03 | -0.09 | 0.11 | 0.022 ± 0.052 |
| 3L | | | | | | | 0.57** | 0.53** | 0.46** | 0.23* | 0.014 ± 0.034 |
| proximal 3R | | | | | | | | 0.96** | 0.91** | 0.17* | 0.034 ± 0.074 |
| In(3R)Mo | | | | | | | | | 0.95** | 0.17* | 0.035 ± 0.083 |
| distal 3R | | | | | | | | | | 0.15* | 0.030 ± 0.072 |
| X | | | | | | | | | | | 0.047 ± 0.006 |

\* 0.0001 < P < 0.05
\*\* P < 0.0001

## DISCUSSION

Our calculations have identified variant (SNP) pairs in 205 lines in the Freeze 2 data set that have linkage disequilibrium (LD) above *r*^2^>0.5. This reveals both the overall patterns of LD, and will facilitate analyses that attempt to disentangle which nucleotides cause phenotypic effects.

Most SNPs in the Drosophila Genome Reference Project are in strong linkage disequilibrium (LD of *r*^*2*^≥0.5) with at least one other variant. More strikingly, many SNPs in the full DGRP with minor allele frequency less than 0.2 are highly correlated with at least one SNP more than 100kbp distant. Thus, while it is true that the DGRP population has low LD relative to other eukaryotes, disequilibrium is still a major element of these data, and careful consideration should be given to its impact at all stages of an association analysis.

Linkage disequilibrium in the DGRP seems to reflect several possible causes (Huang et al. 2014; Pool 2015). First, population-wide LD persists among closely linked SNPs because the recombination events that break down LD are insufficiently rare to counteract the processes such as past admixture, stochastic mutation, drift and potential natural selection. The signature of these events in the DGRP is that local LD remains appreciable throughout the range of SNP frequencies, and is particularly high in regions of reduced recombination. This suggests that local LD would also be found in larger samples of genotypes. The North American population of *D. melanogaster* from which the DGRP lines were sampled was founded by admixture of African and European genotypes (Duchen et al. 2013), and short haplotypes (median 0.17 centiMorgans or 80 kb) of African ancestry can be identified in the DGRP lines (Poll 2015).

Second, variants with low minor allele frequencies (MAF) are on average highly correlated with multiple SNPs throughout the genome. The likely cause of this is random sampling of the very large number of low MAF variants in a relatively small sample of lines. We call this rarity disequilibrium. GWAS analyses often presume that only local LD needs to be considered, but this is not true for variants with MAF less than approximately 0.2 for the DGRP. This source of disequilibrium would be sensitive to the sample size of lines used. Larger samples of genotypes expand the number of possible combinations of lines that would allow a particular minor allele count, and thus confine rarity disequilibrium to a smaller range of MAF. Conversely, smaller samples of subsets of the DGRP lines, would intensify rarity disequilibrium.

Third, the presence of inversions allows differentiation of genotypes carried by each inversion, in turn creating additional LD. The rarity of alternative karyotypes means that this source of LD intensifies the degree of LD already present due to rarity disequilibrium in the DGRP.

Somewhat more speculatively, Pool (2015) has shown that haplotypes inferred to be of African origin in the DGRP have significant positive LD at a genome-wide scale. This finding is difficult to relate quantitatively to our results, as Pool did not report a measure of LD effect size, so it is possible that the degree of LD detected in his study is below the r^2^≥0.5 threshold that we employed. Nevertheless, Pool’s result does suggest that epistatic selection in the Raleigh population could be strong enough to generate LD throughout the genome. For example, if African genotypes at widely spaced genes together enabled an adaptive response to a common environmental challenge, while temperate genotypes at those same loci were favored in the alternative challenge, this could leave a persistent signature of LD. For example, perhaps multi-locus African genotypes are resistant to high temperatures, while multi-locus temperate genotypes are resistant to cold shock. A joint SNP-level analysis of African origin and LD could be very informative.

Similarly, the pattern of correlations in heterozygosity among chromosomal regions we observe (Table 1) suggests that some of the long-distance or inter-chromosomal disequilibrium that we have detected may reflect epistatic selection during the process of inbreeding these lines. Corbett-Detig, et al. (2013) observed that some genotypic combinations in regions distant from each other are observed less frequently than expected in recombinant inbred lines in *D. melanogaster*, as well as other species. The correlations of heterozygosities we detected are the converse of this pattern, but are consistent with selection creating long-distance disequilibrium during the process of inbreeding.

When the karyotypic assignments from three previous studies are in agreement (Huang et al. 2014; Corbett-Detig and Hartl 2012; Langley et al. 2012), our genotype-based assignments of inversion type are concordant, except for three chromosome arms (3R in RAL332, 2R and 3R in RAL409). RAL332 is scored as an inversion heterozygote on the basis of our analysis, and homozygous Standard by Huang et al. and Corbett-Detig and Hartl. This could be due to the loss of the In(3R)Mo after sequencing. The remaining discrepancies between our results and the other scorings cannot be explained on this basis, and are likely either errors in these previous assignments, or perhaps recombination events that have separated karyotype and genotype. Line 409 has two arms previously scored as inverted that have PC scores intermediate between Standard and Inverted types, but no elevated heterozygosity. These arms, plus 2R from RAL 377 could represent recent double recombination events. Similarly the two lines that were both homozygous for arm 2R and assigned an In(2R)NS inversion status in conflict with Corbett-Detig and Hartl’s analysis of inversion breakpoints (RAL69 and 83) each have African ancestry more typical of the alternative karyotype.

Regardless of whether the discrepancies between LD-based genotyping and karyotypic assignments are caused by errors or recombination, our genotypic typing is relevant for those performing association studies, as it summarizes similarity of genotype, and therefore phenotypic effects of the genotypes captured by each inversion. Inversion breakpoints could themselves cause a phenotypic effect, and these are particularly likely to have been involved in the initial spread of rare karyotypes. Coevolution of karyotype and genotype since that time are likely to have altered those initial effects. Our results enable analyses to test whether our LD-based assignments and those based on other criteria give concordant results.

Lines with highly heterozygous regions are potentially more troublesome, as loss of heterozygosity between sequencing and phenotyping cannot be ruled out without additional analysis. This appears to be the most likely explanation for the conflicting evidence concerning the inversion-type of 3R in line RAL332. Association studies should therefore test whether results are affected by inclusion of heterozygous variants and regions which are likely to have diverged in genotype from the sequence data before they are phenotyped.

Our LD calculations will not apply precisely to most association studies based on the DGRP, as each study is likely to use a different subset of lines for phenotyping. We have also calculated the correlations for the 184 lines that we have data for in our own association study (Márquez et al. unpublished). These results show that differences in the genotypes chosen can substantially change the inferred LD structure for rarer SNPs. Nevertheless, the correlations that we have calculated will be useful as the basis for analyses of multi-SNP associations. For example, after identifying a set of SNPs with significant associations, one could reanalyze those in multi-SNP analyses that include the most highly correlated SNPs to diagnose which SNPs are most likely to represent the variants that cause phenotypic differences, and which have their signal confounded with those from other SNPs. If such SNPs are all nearby, the inference of that genomic region as causal can be strong, even if the precise nucleotide responsible remains unknown. Follow-up studies of such regions are likely to be worthwhile. In contrast, SNPs for which the addition of distant SNPs renders effects ambiguous would be poor candidates for follow-up studies.

## ACKNOWLEDGEMENTS

This work was supported by NIH 1R01GM094424-01. We thank two anonymous reviewers for thoughtful comments. Computing support was provided by the Research Computing Center at Florida State University.

